# Spatial flexibility of distractor suppression in the tactile modality

**DOI:** 10.1101/2025.03.10.642407

**Authors:** Alyssa S. Ignaco, Nicole Serino, Alexa Trzpis, Edmund C. Lalor, Manuel Gomez-Ramirez

## Abstract

Object sensing and manipulation rely on the somatosensory system’s ability to relay relevant information about a skin location encoding tactile information about the object while suppressing irrelevant or distracting inputs on the skin—a process known as distractor suppression. Although tactile selective attention is flexible, capable of being allocated to non-contiguous skin areas on the hand and with spatial resolution limited to within a finger, the spatial properties of distractor suppression in touch remain poorly understood. In particular, studies have not explored the effects of proactively cueing distractor locations in touch, leaving open questions about whether active deployment of distractor suppression operates independently of attentional enhancement. To address this gap in knowledge, in this study, participants performed an amplitude discrimination task on the hand in the presence of distractors that were cued in advance. Our data revealed that validly cueing distractor locations improves discrimination accuracy and reduces reaction times of behaviorally-relevant tactile stimuli. The data also showed that behavior is unaffected when attended stimuli are flanked by distractors within or across fingers, suggesting that distractor suppression mechanisms can be split across different skin locations on the hand. These findings provide novel insights into the spatial allocation and flexibility of distractor suppression mechanisms in touch, and underscore the importance of proactive cueing in shaping these mechanisms.

## INTRODUCTION

When manipulating an object with the hand (haptics), our fingers work together to derive a representation of the object that is critical to grasping the object. The sense of touch plays a key role in haptics by providing relevant sensory signals to motor planning areas that mediate grasping behavior. A major function of the somatosensory system is to relay relevant information about the target object, while ignoring all distracting tactile information impinging on the hand (Hsiao & Gomez-Ramirez, 2012). Consider the following example: When searching for keys in a bag without the help of visual cues, it is advantageous to selectively process touch information on the fingertips and ignore all other information impinging on other locations on the hand (e.g., the back of the hand). Selective attention plays an important role in these situations by enhancing information relevant to our haptic goal while at the same time suppressing distractor cues (Adler et al., 2009; Evans & Craig, 1991; Frings et al., 2008; Gomez-Ramirez et al., 2016a). The ability to ignore irrelevant sensory information is mediated by an active mechanism called distractor suppression (Gaspelin et al., 2015; Geng, 2014; Noonan et al., 2016; Schneider et al., 2022), a functional mechanism that is prevalent in other sensory modalities (Braithwaite et al., 2010; Chang & Egeth, 2019; Hughes, 2014; Schneider et al., 2022; Schwartz & David, 2018). Tactile distractor suppression is crucial to haptics by ensuring that the limited resources of the brain are not allocated to processing irrelevant sensory information, providing preferential access of relevant touch inputs across the sensori-motor circuit that facilitates haptics. To date, most studies on distractor suppression have been performed with cues that direct attention to the relevant stimuli only. Researchers then analyze activity in areas where distractors may occur to determine how the brain suppresses irrelevant information (i.e., in locations not predicted by the attention cue) (Ede et al., 2011; Eimer & Forster, 2003b; Forster & Eimer, 2005; J. J. Foxe & Snyder, 2011; Gomez-Ramirez et al., 2016a; Haegens et al., 2012a; Jones et al., 2010; Kelly et al., 2008, 2010; Mohr et al., 2020; Sacchet et al., 2015; van Ede et al., 2012, 2014a). Because of this experimental manipulation, some studies have questioned whether distractor suppression is separable from mechanisms that enhance attended stimuli (Gaspelin & Luck, 2018; Noonan et al., 2018; van Moorselaar & Slagter, 2020). To address this gap in knowledge, we conducted an experiment that proactively cued the location of distractors to study whether distractor suppression enhances discrimination of behaviorally-relevant tactile stimuli presented to the hand. We hypothesize that if distractor suppression operates independently from attentional enhancement, performance should be enhanced when distractors are valid vs. invalidly cued while the attended location remains the same.

Selective attention mechanisms are highly flexible across multiple dimensions, such as space and object features (Gomez-Ramirez et al., 2016b; Moore & Zirnsak, 2017; Snyder & Foxe, 2010). Many studies have metaphorically described attention as acting as a “spotlight”, enhancing sensory processing in brain areas that represent the attended locations. In particular, studies in vision have shown that selective attention can be split across discrete retinotopic areas encoding the location of the relevant stimuli, as opposed to acting as a unitary spotlight within an area (Jans et al., 2010; McMains & Somers, 2004; Vanrullen & Dubois, 2011). Similar observations have been made for suppression of distracting visual stimuli (Drisdelle & Eimer, 2023; Frey et al., 2014). The study by Frey et al. (2014) focused on whether the alpha suppression effect (J. Foxe & Snyder, 2011), observed using electroencephalography (EEG), follows the single vs. divided spotlight hypothesis. Topographic analyses of alpha-band (8-14Hz) activity, a neural correlate of active suppression effects (J. J. Foxe et al., 1998; Haegens et al., 2012b; van Ede et al., 2014b; Worden et al., 2000), demonstrated multiple foci of increased alpha in the occipito-parietal area when task demands required ignoring multiple non-contiguous spatial locations, a finding that supports the divided spotlight hypothesis (Frey et al., 2014).

Similar to visual studies (Frey et al., 2014; Jans et al., 2010; McMains & Somers, 2004), work in the somatosensory modality show that tactile selective attention can be allocated to different body locations simultaneously (Clepper et al., 2021; Craig, 1985; Daniel et al., 2022; Eimer & Forster, 2003a; Forster et al., 2016; Gherri et al., 2023). In particular, EEG data showed that when attending to two nonadjacent fingers on one hand (e.g., the index and little finger), neural responses to unattended intervening fingers (e.g., the middle and ring finger) are reduced (Eimer & Forster, 2003a). These results indicate that tactile attention can be allocated to non-contiguous areas within the same hand.

In the present study, we investigated whether active mechanisms of distractor suppression in touch can also be flexibly allocated to non-contiguous areas on the hand. In two different experiments, human participants performed a vibrotactile amplitude discrimination task on the hand with distractor stimuli presented in different spatial arrangements relative to attended stimuli. In Experiment 1, we test whether suppression can be deployed when distractor locations are flexibly cued on a trial-by-trial basis. In Experiment 2, we test whether flexibly cueing attended vs. distractor stimuli is beneficial for discrimination of the attended stimulus. We found that participants’ performance is enhanced when distractors are validly cued, and that invalidly cued distractors shift participants’ perceptual thresholds. These observations suggest that proactive deployment of distractor suppression is beneficial to perception in similar ways as mechanisms of attentional enhancement. The data also show that participants’ behavior is unaffected when attended stimuli are flanked by distractors cued within or across fingers, suggesting that distractor suppression mechanisms can be split across different skin locations on the hand. Taken together, our behavioral findings shed light on the mechanisms that underlie the spatial deployment of selective attention and distractor suppression in the tactile modality.

## METHODS

### Participants

Thirteen healthy, right-handed individuals were recruited for Experiment 1 (9 females and 4 males, mean ± SD age = 22.7 ± 2.50 years old). Five participants completed one experimental session in one day, while eight participants completed two experimental sessions held on two different days. Twenty-seven healthy individuals were recruited for Experiment 2 (17 female and 10 males, mean ± SD age = 21.8 ± 5.58 years old). Data of three participants from Experiment 1 and seven participants from Experiment 2 were discarded because they had strong biased responses that flattened their psychometric curves or very low number of trials in some conditions (see Data Analysis section for a detailed description). Participants were recruited via the University of Rochester’s email lists and informal referrals. All participants gave written informed consent and were monetarily compensated for their time. The study procedures were approved by the Institutional Review Board (IRB) at the University of Rochester.

### Tactile Stimuli and Experimental Setup

Participants’ left hand was secured on a custom-made exoskeleton (see **Figure 1A**) that used finger splints to accommodate and secure the fingers. Movements of the stimulated fingers were additionally restricted by applying a small amount of nontoxic adhesive to the fingernails of the index, middle, and ring fingers (i.e., digits 2, 3, and 4). Vibrotactile stimulation was provided using piezoelectric bimorph bending actuators (Model #BA4010, PiezoDrive, https://www.piezodrive.com/; see inset in **Figure 1A**). The piezo actuators were affixed to mechanical structures designed to securely hold the devices in place. Rounded and custom-made 3D-printed plastic attachments (diameter = 5 mm) were placed at the ends of the piezo actuators that were placed on different finger pads of digits 2, 3, and 4 (D2, D3, and D4, respectively) of the left hand (see **Figure 1A**). A National Instruments Data Acquisition System (NI-DAQ) controlled by custom-written MATLAB code was used to control vibrotactile stimulation.

**Figure 1.**
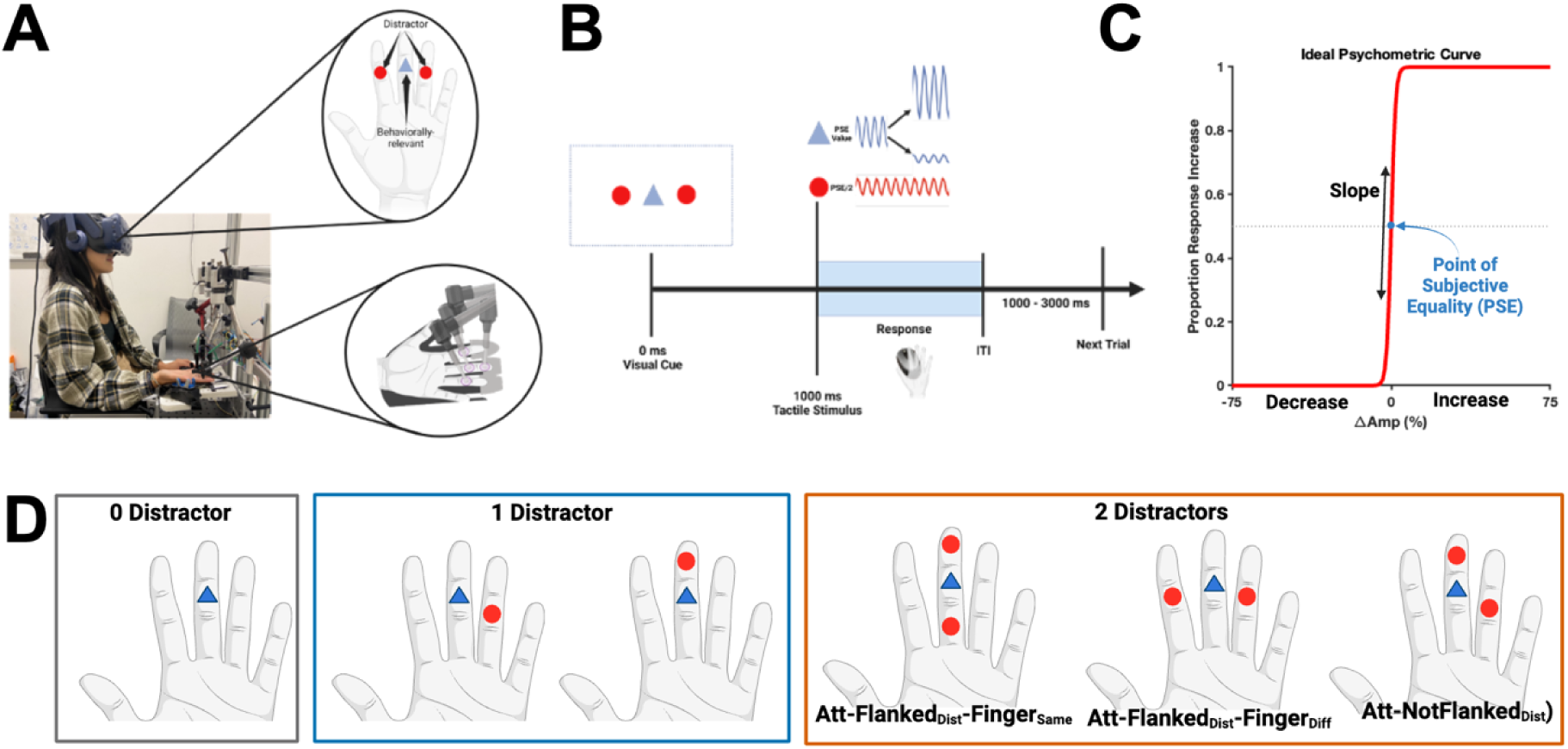
Experimental setup and trial timeline. (**A**) Participants sat with their left hand supinated (palm up) resting on top of a padded hand rest. Participants wore a VR headset to provide visual cues to the targets and distractors, and block all visual information related to the tactile stimuli. Pink auditory noise was presented via built-in headphones on the VR to block out noise from the tactile stimulators. Four vibrotactile stimulators were placed on their finger pads (middle pad of D2, distal and middle pads of D3, and proximal pad of D3 or middle pad of D4). The picture is from a co-author in the study, and was taken during the experimental piloting. (**B**) Each trial began with a visual cue showing a blue triangle for the target location and red circle(s) for the distractor location(s). After 1000 ms, the target and distractor vibrations were provided simultaneously. The target vibration started at a baseline amplitude, determined by each subject’s point of subjective equality (PSE, or perceptual threshold), then increased or decreased in amplitude after 500 ms for another 500 ms. The distractor vibrations remained at a constant amplitude, which was set at half of the baseline amplitude of the target vibration. Participants were asked to make their response via a mouse click as soon as they felt confident about their selection. **(C)** Ideal psychometric curve. Sensitivity refers to the slope, or steepness, of the curve, while the point of subjective equality (PSE) is the amplitude change at which participants’ responses fell to chance level (i.e., 50%). The PSE would ideally be at 0%, or no change in amplitude. **(D)** Illustration of representative examples of the different distractor arrangements for each distractor number condition. Att-Flanked_Dist_-Finger_Same_ = Attended Stimulus Flanked by Distractors on Same Finger, Att-Flanked_Dist_-Finger_Diff_ = Attended Stimulus Flanked by Distractors on Different Finger, and Att-NotFlanked_Dist_ = Attended Stimulus Not Flanked by Distractors.

The vibration stimulus (y) consisted of a sine wave created from Equation 1:

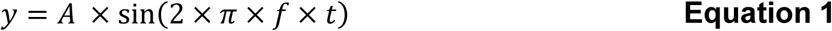

Where *A* represents the amplitude of the stimulus that was adjusted to manipulate the vibration intensity. *F* represents the frequency of the sine wave, which remained constant at 200 Hz for all stimuli, and *t* is the duration of the stimulus (1000 ms). The latter half of the tactile stimulus (i.e., 500 to 1000 ms) had a change in amplitude, relative to the first half of the stimulus (see e.g., **Figure 1B** *inset blue traces*), that varied across seven levels relative to each participant’s point of subjective equality (PSE) for amplitude discrimination (−75%, −25%, −15%, 0%, 15%, 25%, 75%). For an ideal observer, 0% represents the PSE (or perceptual threshold, **Figure 1C**). However, because human participants have different perceptual biases, we determined each individual’s PSE value in a session prior to the experimental task (see below for details). Participants wore a virtual reality (VR) headset (HTC VIVE Pro) to present visual stimuli and limit visual cues from the tactile stimulators. Visual cues were presented using the Unity Real-Time software. We delivered ‘pink noise’ stimuli through the VR’s headphones to mask sounds coming from the bending movement of the piezo actuators. Custom code written in MATLAB was used to control the timing of the tactile, visual, and auditory stimuli.

### Pre-Experimental Task to Determine the PSE of Participants

We first established how sensitive and biased each participant was to detecting amplitude changes to adjust the intensity of the attended stimuli to their own sensitivity. To do this, we established a perceptual threshold for each participant where their ability to detect an amplitude change in the stimulus fell to chance level (i.e., 50%). This perceptual threshold is also known as the PSE.

To determine the PSE in Experiment 1, participants performed an amplitude discrimination task (N = 56 trials), in the absence of distractors, that required them to judge whether the vibratory stimulus on the middle pad of D3 increased or decreased in intensity (see Experimental Task section for details of the task). The PSE_PreTask_ value was determined as the amplitude change that yielded a participant to equally respond that the stimulus increased or decreased in intensity (i.e., a 50% choice response across trials).

To determine the PSE in Experiment 2, participants received a stimulus on the middle pads of D2, D3, D4, or the distal and proximal pads of D3. Thus, we determined participants’ PSE_PreTask_ at each of these skin locations by performing a stepwise amplitude discrimination task, where the vibrotactile stimulus started off with a large increase (75%) in amplitude then gradually decreased until the participant’s responses began to fluctuate between decrease and increase (i.e., 50% responses). The amplitude change of a trial would decrease or increase by 10% of the preceding trial’s amplitude change depending on whether the participant responded correctly (e.g. if the amplitude change was an increase, the following trial’s amplitude change decreased or increase by 10% of the prior amplitude change if the participant responded correctly or incorrectly, respectively). There was also a 25% chance that the amplitude of a trial randomly decreased by 75% (i.e., - 75%) to ensure that participants remained attentive during the task, and separate long stretches of trials with increases in amplitude. For every trial following the tenth trial, linear regression analysis was performed on the amplitude changes using the previous 10 trials (excluding the 25% of trials that randomly decreased by −75%) until the resulting slope was not significantly different from 0 (p > 0.05). After achieving a non-significant linear slope, the amplitude change of the last trial was considered to be the PSE_PreTask_ at the stimulus location.

### Attention Experimental Tasks

#### Experiment 1

A trial started with the presentation of a blue triangle and red circles on a visual monitor to cue the locations of the attended and distractor stimuli, respectively (**Figure 1B**). Following 1000 ms the onset of the visual cue, tactile stimuli were delivered to the hand. Participants were instructed to judge whether there was an increase or decrease in the amplitude of the vibratory stimulus in the attended finger location only. Participants made a left or right mouse button push if they perceived that the stimulus decreased or increased in amplitude, respectively. Following the response, an inter-trial interval between 1000 and 3000 ms was presented (uniformly randomized). Distractor stimuli were presented at a constant vibration amplitude of half the intensity relative to the baseline amplitude of the attended stimulus. Participants were instructed to ignore all stimuli presented in the cued distractor locations.

**Figure 1D** shows different spatial arrangements of attended and distractor stimuli presented to the hand. The location of the attended stimulus was fixed to the middle pad of D3. Distractor stimuli were presented on the same finger (e.g., proximal or distal pads of D3), or different fingers (e.g., the middle pads of D2 and D4) in relation to the attended stimulus. The same-finger and different-finger distractor/attended stimuli arrangements were presented in separate behavioral sessions due to experimental constraints related to the placement of the piezo motors. The order of the same-finger vs. different-finger stimuli arrangement blocks were randomized across participants. A Wilcoxon signed-rank test did no show a difference in performance between the first and second behavioral sessions (W = 13, p = 0.5469). For each behavioral session, distractors could occur in any combination of three possible locations (i.e., Different-finger session – middle pads of D2 and D4 and distal pad of D3; Same-finger session – middle pad of D2 and distal and proximal pads of D3). Attended stimuli were presented alone (i.e., no distractors) or in the presence of one or two distractors. Trials with two distractors were presented in flanking or non-flanking spatial arrangements relative to the attended stimulus. In the same-finger session, distractors in a flanking arrangement were located on the distal and proximal pads of D3. In the different-finger session, distractors in a flanking arrangement were located on the middle pads of D2 and D4. Distractors in the non-flanking arrangement were located on the distal or proximal pads of D3 and the middles pads of D2 or D4. That is, one distractor was presented on D3, and the other distractor was presented on a neighboring finger of D3. On 75% of the trials, the distractor(s) were presented on the same location predicted by the visual cue (validly cued distractor), while on the remaining 25% of trials, distractors were randomly presented to the other finger locations that a distractor could be presented (invalidly cued distractor). The different distractor conditions were randomly presented on each trial. Participants performed a total of 756 trials in each experimental session (63 trials per block, 12 blocks of trials), resulting in 12-18 trials for each experimental condition in each stimulus amplitude level.

#### Experiment 2

Experiment 1 tested whether suppression could be flexibly allocated to distractor locations on a trial-by-trial basis. In Experiment 2, we tested whether cueing the attended vs. distractor locations was more beneficial for discrimination (see **Supplemental Figure 1**). For each cue type (attended and distractor), one location had a ring around the cue to indicate that there was a high probability that the stimulus would be presented in that location (i.e., a valid stimulus). The cued location without the ring represented a low-probability location (i.e., an invalid stimulus). On separate blocks, we varied the location of the validly- and invalidly-cued stimuli for one of the cue types while keeping the locations for the other cue type fixed (e.g., presenting the cues for validly- and invalidly-cued attention stimuli in different finger locations, while keeping the validly- and invalidly-cued distractor stimuli on the same locations for the entire block). The order of the fixed cue type was alternated across blocks and randomized across participants. The cue locations were designed to match the two-distractor flanking and non-flanking arrangements from Experiment 1. Participants performed the same amplitude discrimination task as in Experiment 1. On each trial, the participant received one attended stimulus and one distractor stimulus simultaneously 1000 ms after visual cue onset. Eleven of the 27 participants also received no distractor on 30% of the trials, randomly. The attended and distractor stimuli occurred at their valid locations on 75% of the trials and at their invalid locations on 25% of the trials. The experiment was performed in a single session. The participants who did not have the No Distractor condition performed a total of 600 trials (24 blocks of 25 trials each), while participants who had the No Distractor condition performed a total of 900 trials (30 blocks of 30 trials each). The amplitude change of the attended stimulus was randomly chosen per trial with weights of 8%, 17%, 21%, 8%, 21%, 17%, and 8% for the −75%, −25%, −15%, 0%, 15%, 25%, and 75% amplitude change condition, respectively.

### Data Analysis

Even though the PSE of each participant was determined in the pre-experimental task, some participants still showed a strong bias in reporting an increase or decrease in stimulus amplitude during the attention experimental task. These participants were identified and excluded from the analysis. Their overall psychometric curves are shown in **Supplemental Figure 2**. To identify these participants, we performed permutation tests for each stimulus amplitude change condition to determine the proportion of times that a participant reported an increase or decrease in amplitude change (N = 20% of trials for each amplitude change condition, repeated 5000 times). If the proportion value was above or below 0.5 on more than 95% of all trials for at least five amplitude change level conditions, then we removed the participant from analyses. Two additional participants were excluded from Experiment 2 because of low number of trials in the flanking conditions.

Using the data from the remaining participants, we derived cross-subject psychometric curves by computing the median of the mean responses across those participants for each stimulus amplitude level condition. We derive cross-subject psychometric curves rather than deriving each subject’s psychometric curve because we did not have the power (i.e., number of trials) to fit psychometric functions at the individual participant level. A Logistic function was fitted to determine the slope and PSE values of the psychometric curves (Equation 2):

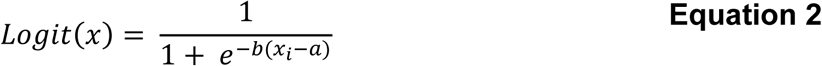

Where *x_i_* denotes the vibration amplitude change level, *a* is the 50% mark of the distribution (i.e., PSE value), and *b* is the slope of the psychometric function (i.e., sensitivity). An ideal psychometric curve is shown in **Figure 1C**. Statistical effects on the parameters of the psychometric curve were estimated using permutation tests, with a significance threshold below the 0.05 level. First, we built a surrogate distribution for each parameter of the model using bootstrapping techniques (N = 1000 iterations). On each iteration, we fitted **Equation 2** using 30 trials per stimulus amplitude level (N = 210 trials per iteration) that were randomly chosen with replacement from the pooled choice responses across all participants. Surrogate distributions were derived for each experimental condition (i.e., Cue Validity, Distractor Number, and Distractor Spatial Arrangement). To test how Cue Validity (Valid vs. Invalid distractor cue) modulates a parameter of the model, we pooled the bootstrapped distributions for the model’s parameter of interest (i.e. *a* or *b*) across the Valid and Invalid conditions. After pooling the data, we computed the difference in means between two randomly-sampled sub-distributions (composed of 30 samples each; N = 5000 times). We compared the number of times that the difference-distribution (i.e., the 5000-sample difference distribution) was greater than the difference between the means of the surrogate distributions of the Valid and Invalid conditions. A value below 5% was classified as a statistically significant effect. The same shuffling analyses were performed to test how Distractor Spatial Arrangement (Attended Stimulus Flanked by Distractors on Same Finger, Attended Stimulus Flanked by Distractors on Different Fingers, and Attended Stimulus Not Flanked by Distractors) modulates PSE (i.e., the parameter ‘*a’* of the model) and discrimination sensitivity (i.e., the parameter ‘*b’* of the model).

To test how the number of distractors modulates PSE and/or perceptual sensitivity, we randomly selected 30 PSE or sensitivity values from the surrogate bootstrapped distribution (see previous paragraph) of each Distractor Number condition (0, 1, and 2 distractors). Afterwards, we performed a linear regression analysis using Distractor Number as a predictor and calculated the slope value. This regression analysis was repeated 5000 times. We then combined and shuffled the PSE or sensitivity data across the three Distractor Number surrogate bootstrapped distributions, randomly assigned 30 datapoints to each Distractor Number level (0, 1, and 2), and performed linear regression analysis on the randomly selected data. We performed 5000 iterations of the linear regression analysis on the shuffled data to obtain a distribution of linear slopes. To test for statistical effects, we compared the number of times that the shuffled-slope distribution was greater than the mean of the slope distribution computed from the non-shuffled Distractor Number conditions. A value less 5% was classified as a statistically significant effect.

Statistical effects on behavioral accuracy (i.e., Hits) and reaction time (RT) were calculated using a generalized linear mixed-effects model (GLME) with an inverse gaussian distribution. The factors of GLME were Distractor Number, Cue Validity, Distractor Arrangement, or Stimulus Amplitude Level. Pairwise Wilcoxon signed-rank tests were then performed for predictor variables with significant p-values (p < 0.05) from the GLME model.

We assessed whether participants’ individual response biases modulate the effects that distractors have on behavioral performance (e.g., larger distractor interference effects in participants with lower PSEs). For each distractor number condition, we performed separate linear regression analyses using participants’ PSE_PreTask_ as a dependent variable (determined from the ‘Experimental Task to Determine the PSE of Participants’) and accuracy or RT as the dependent variables.

## RESULTS

### Effects of number of distractors on behavioral performance

Vibration discrimination is impaired as a function of distractor number (**Figure 2A**). The psychometric curves (*left graph* in **Figure 2A**) demonstrate a leftward shift in perceptual thresholds (PSEs; 50% threshold of the psychometric curve) as the number of distractors increases. Non-parametric shuffling tests show that the PSEs across Distractor Number condition are significantly different from each other, with a slope value of −10.97 (p = 0; see middle *upper graph* **Figure 2A**). Distractor Number also has an effect on perceptual sensitivity (*left graph* in **Figure 2A**), with sensitivity values decreasing with a linear slope value of −1.58 (p = 0; *middle lower graph* **Figure 2A**). RTs are also slower in the presence of distractors (*right graph* in **Figure 2A**). A GLME on RTs with Stimulus Amplitude Level and Distractor Number as factors demonstrated a significant effect of Distractor Number (F(1,206) = 7.1363, p = 0.0082), but no significant effects for Stimulus Amplitude Level (F(1,206) = 0.7725, p = 0.3805) nor an interaction effect (F(1,206) = 1.9139, p = 0.1680). Taken together, these data show that tactile distractors interfere with participants’ ability to discriminate changes in vibrations, with interference effects increasing with the number of distractors.

**Figure 2.**
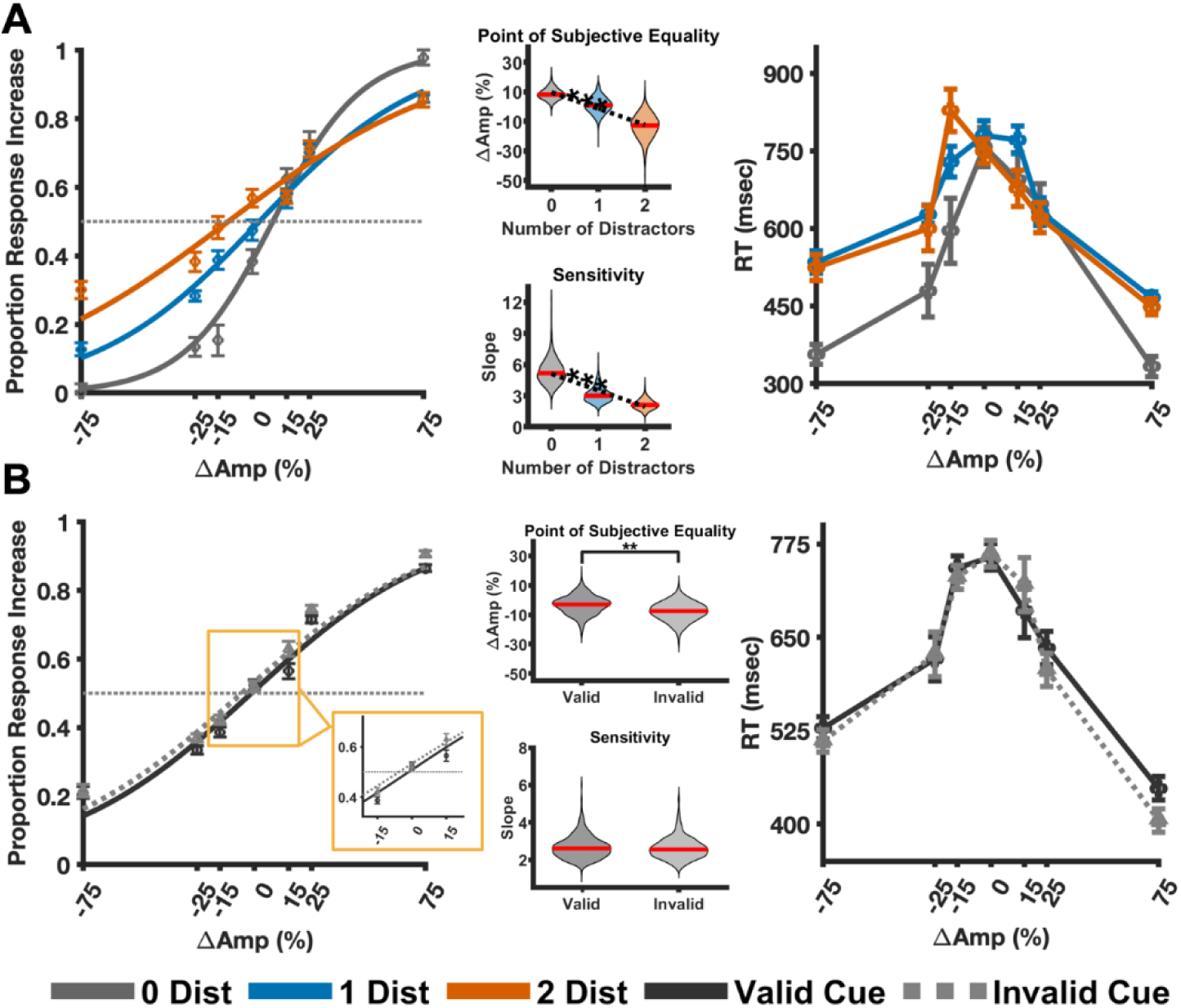
Distractor and cueing effects on perceptual sensitivity, response bias, and reaction time. (**A**) The left graph shows the psychometric functions for each distractor number condition. The solid lines are curve fits to the human data using a logistic function. The curve fits produced R^2^ values of 0.83, 0.75, and 0.66 for the 0, 1, and 2 Distractor conditions, respectively. Violin plots of the psychometric fit parameters, Point of subjective equality (PSE, middle-top) and Perceptual Sensitivity (middle-bottom). The solid red lines are the median values of each distribution, and the dotted black lines are the linear fits as a function of number of distractors. The right graph shows the median RTs for each stimulus amplitude level. (**B**) Same plots as (A) but for the Valid (solid black) vs. Invalid cue (dotted gray) distractor conditions collapsed across number of distractors. The R^2^ for the validly and invalidly cued conditions are 0.76, and 0.70, respectively. * = p < 0.05, ** = p < 0.01, *** = p < 0.001. N = 10. Error bars indicate ± within-subject standard error of the mean (SEM).

### Effects of distractor cueing on behavioral performance

PSEs are shifted by invalid cueing of the distractor stimuli (**Figure 2B**). Non-parametric shuffling tests show that the PSEs between validly- and invalidly-cued distractor stimuli are significantly different from each other (p = 0.0050; *upper middle panel* **Figure 2B**). However, distractor cue validity did not have a significant effect on perceptual sensitivity (p = 0.3714; *lower middle panel* **Figure 2B**). A GLME on RTs with factors of Cue Validity (Valid vs. Invalid) and Stimulus Amplitude Level did not reveal differences between Cue Validity (F(1,136) = 0.0112, p = 0.9159, *right panel* **Figure 2B**), or Stimulus Amplitude Level (F(1,136) = 1.519, p = 0.2199). The GLME also did not show a significant interaction effect between Cue Validity and Stimulus Amplitude Level (F(1,136) = 0.1519, p = 0.6973).

Our experimental paradigm makes specific predictions as to how distractors affect discrimination of stimuli with an increase vs. decrease change in amplitude. Specifically, the added sensory input from distractors may ‘mask’ (and even reverse) the perception of an amplitude decrease that would lead to a reduction in correct behavior. In contrast, for stimuli with amplitude increases, the added sensory input from the distractor(s) may have beneficial effects in behavior. To test this hypothesis, we separately analyzed the effects of Distractor Number and Cue Validity on accuracy and RT for stimuli with amplitude decreases (−75%. −25%, −15%) and amplitude increases (15%, 25%, 75%). A GLME test for stimuli with amplitude decreases revealed a main effect of Distractor Number (0, 1, or 2 Distractors) on behavioral accuracy (F(1,56) = 7.7990, p = 0.0071; *left graph* in **Figure 3A**). Pairwise Wilcoxon signed-rank tests between Distractor Number conditions revealed significant differences in accuracy between 0 vs. 2 distractor stimuli (W = 55, p = 0.002), and between 1 vs. 2 distractor stimuli (W = 55, p = 0.002), but no differences between 0 vs. 1 distractor stimuli conditions (W = 40, p = 0.2324). The GLME test also revealed a main effect of Cue Validity (F(1,56) = 10.6648, p = 0.0019), with enhanced behavior for validly- vs. invalidly-cued distractor stimuli (*solid vs. dashed bars in the left graph* of **Figure 3A**). The GLME test also revealed an interaction effect between Distractor Number and Cue Validity (F(1,56) = 5.0982, p = 0.0279). Pairwise Wilcoxon signed-rank tests showed a significant difference in accuracy between validly- vs. invalidly-cued distractor stimuli for the 0 Distractor condition (W = 52, p = 0.0098) but not for 1 Distractor (W = 29, p = 0.9219) or 2 Distractors (W = 26, p = 0.9219). A GLME test for stimuli with amplitude increases did not show effects of Distractor Number on behavioral accuracy (F(1,56) = 0.3298, p = 0.5681) nor Cue Validity (F(1,56) = 0.0015, p = 9697). The GLME also did not reveal an interaction effect between Distractor Number and Cue Validity (F(1,56) = 0.0224, p = 0.8815; *right bar plots* in **Figure 3A**).

**Figure 3.**
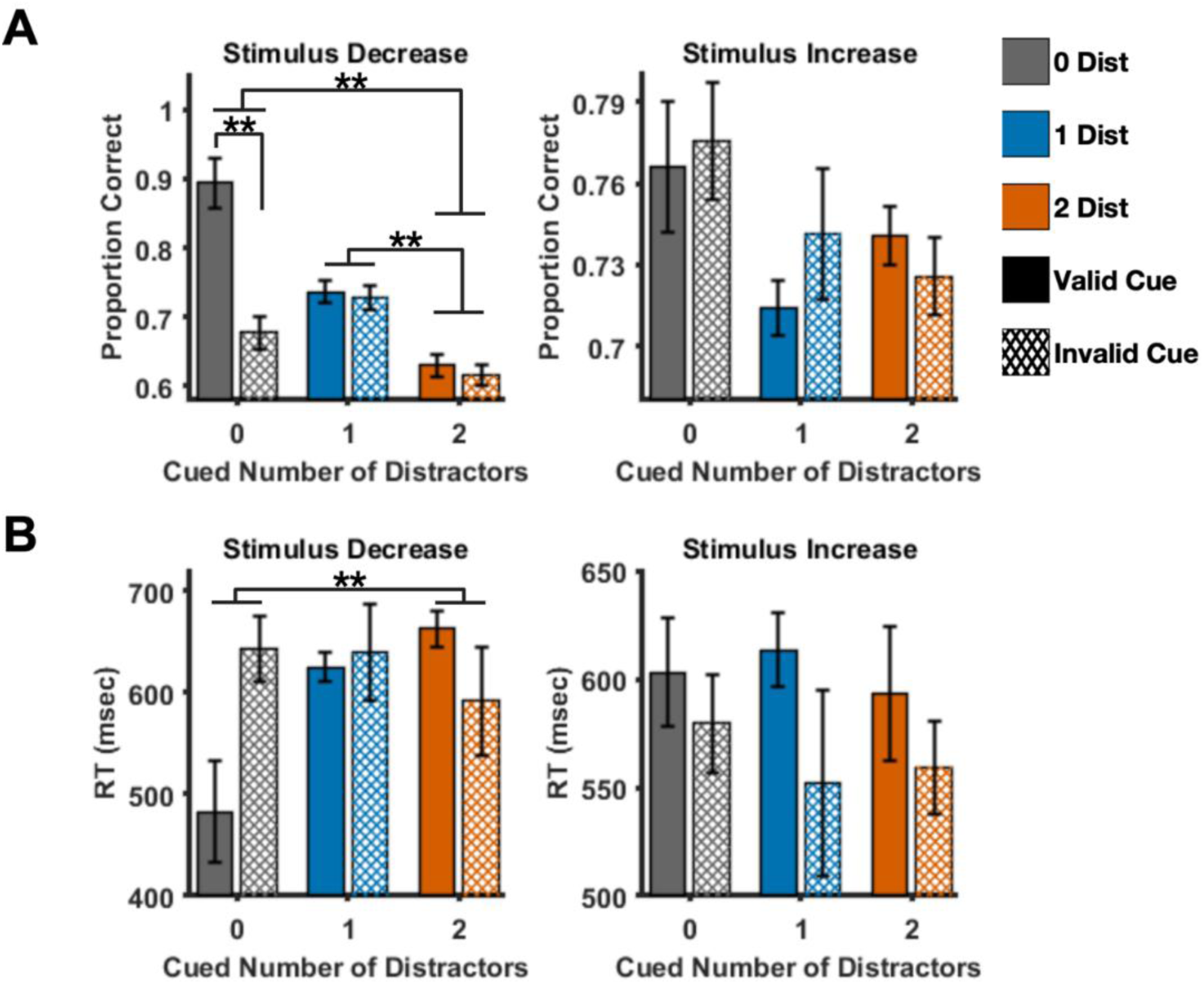
Distractor and cueing effects based on stimulus amplitude change direction. (**A**) The left and right graphs show the median accuracy across participants for stimuli with a decrease and increase amplitude change, respectively. (**B**) The left and right graphs show the median RT across participants for stimuli with a decrease and increase amplitude change, respectively. * = p < 0.05, ** = p < 0.01. N = 10. Error bars indicate ± within-subject SEM.

The number of distractors systematically increases the RT of stimuli with decreasing amplitude. A GLME model on RTs for stimuli with an amplitude decrease revealed a main effect of Distractor Number (F(1,56) = 4.2198, p = 0.0446). Post-hoc Wilcoxon signed-rank tests only showed significant differences in RT between 0 vs. 2 distractor stimuli (W = 8, p = 0.0488) but not for 0 vs. 1 Distractor (W = 9, p = 0.0645) nor 1 vs. 2 Distractor stimuli (W = 19, p = 0.4316). The GLME also revealed a trend towards significance for Cue Validity (F(1,56) = 2.7599, p = 0.1022), with slightly faster RTs for validly- vs. invalidly-cued RTs. However, the GLME model did not show an interaction effect between Cue Validity and Distractor Number (F(1,56) = 1.8714, p = 0.1768). A GLME test on stimuli with an amplitude increase revealed no effects of Distractor Number (F(1,56) = 0.1675, p = 0.6839) nor Cue Validity (F(1,56) = 0.0142, p = 0.9056) on RTs. The GLME also did not show an interaction effect (F(1,56) = 0.1489, p = 0.7011). Taken together, these results suggest that distractor effects modulate according to the amplitude change of the attended stimulus.

### Relationship between distractor interference effects and subject response bias

Participants with a bias for responding that a stimulus increased in amplitude (i.e., negative PSEs) demonstrate greater distractor interference effects for stimuli that decrease in amplitude. We performed linear regression analyses on the differences in performance between the distractor (1 or 2 Distractor) and 0 Distractor conditions to compare the distractor interference effects as a function of participants’ PSE_PreTask_ (i.e., subtracting accuracy in the 0 Distractor condition from the 1 or 2 distractor conditions). The linear regression analyses revealed a significant relationship for 1 Distractor (t(8) = 3.9904, p = 0.0040, R^2^ = 0.6656; *blue line on the left graph* of **Figure 4A**) and 2 Distractor conditions (t(8) = 2.9849, p = 0.0175, R^2^ = 0.5269; *orange line on the left graph* of **Figure 4A**) for stimuli that decrease in amplitude. For stimuli that increase in amplitude, linear regression analyses did not reveal significant effects for any Number of Distractor condition (*right graph* of **Figure 4A**). Linear regression analyses on the RT revealed a significant effect in the 1 Distractor condition (t(8) = −3.0396, p = 0.01061, R^2^ = 0.5359; *blue line on the left graph* of **Figure 4B**) and a trend towards significance in the 2 Distractor condition for stimuli that decrease in amplitude (t(8) = −2.1244, p = 0.0664, R^2^ = 0.3607; *orange line on the left graph* of **Figure 4B**). Linear regression analyses showed no significant relationships between RT and PSE_PreTask_ (*right graph* of **Figure 4B**). These results indicate that distractor interference effects are enhanced for individuals with response biases for stimuli that decrease in amplitude.

**Figure 4.**
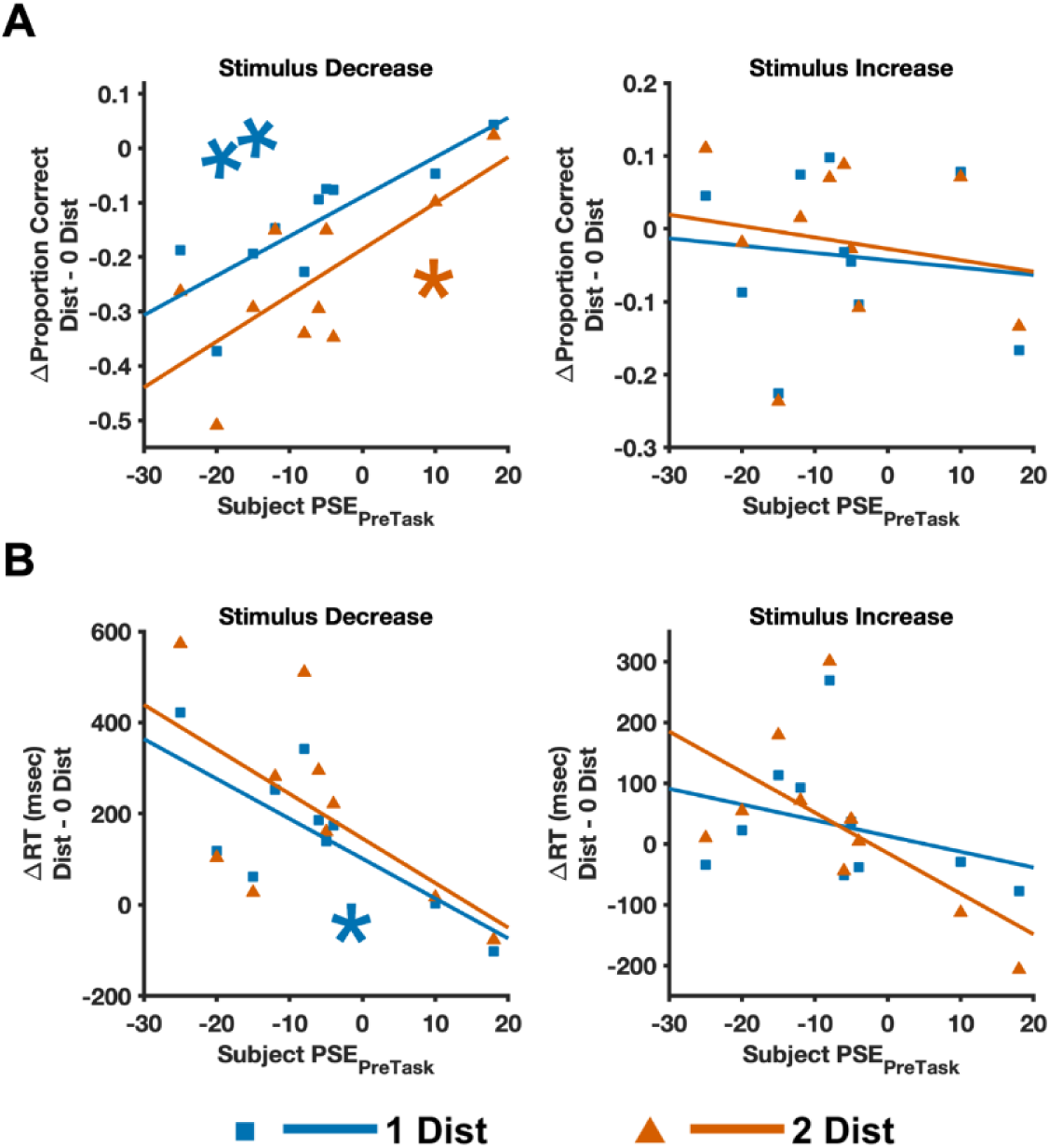
Effects of participants’ biases on distractor interference effects. (**A**) Relationship between performance accuracy and PSE_PreTask_. Each data point represents the accuracy difference between the distractor (1 or 2 distractors) and the 0 distractor conditions of each participant for the 1-distractor (blue squares) and 2-distractor (orange triangles) conditions. The solid lines represent the linear regression plots between the accuracy differences and PSE for stimuli with decreased amplitudes (left graph) and stimuli with increased amplitudes (right graphs). **(B)** Same as (A) but for a relationship between RT and PSE_PreTask_. N = 10. * = p < 0.05, ** = p < 0.01

### Effects of distractor spatial arrangement on performance

Distractor interference effects depend on the spatial arrangement of distractor stimuli. **Figure 5A** shows psychometric curves for 2-Distractor stimuli conditions arranged in a flanking pattern within one finger (Attended Stimulus Flanked by Distractors on the Same Finger, Att-Flanked_Dist_-Finger_Same_; magenta trace), flanking pattern across fingers (Attended Stimulus Flanked by Distractors on Different Fingers, Att-Flanked_Dist_-Finger_Diff_; green trace), or non-flanking pattern across fingers (Attended Stimulus Not Flanked by Distractors, Att-NotFlanked_Dist_; orange trace). Pairwise shuffling statistics tests revealed significant differences in PSE between Att-Flanked_Dist_-Finger_Same_vs. Att-Flanked_Dist_-Finger_Diff_ (p = 0), Att-Flanked_Dist_-Finger_Same_ vs. Att-NotFlanked_Dist_ (p = 0), and Att-NotFlanked_Dist_vs. Att-Flanked_Dist_-Finger_Diff_ (p = 0), with the Att-Flanked_Dist_-Finger_Same_and Att-Flanked_Dist_-Finger_Diff_ conditions having the lowest and highest PSE values, respectively (*upper panel* **Figure 5B**). Pairwise shuffling statistics tests also revealed significant differences in perceptual sensitivity between Att-Flanked_Dist_-Finger_Same_vs. Att-Flanked_Dist_-Finger_Diff_ (p = 0.0214), and Att-NotFlanked_Dist_ vs. Att-Flanked_Dist_-Finger_Diff_ conditions (p = 0.0104; *lower panel* **Figure 5B**). These data suggest that changes in vibration amplitude are easier to discriminate when there are no distractors present within the same finger as the attended stimulus.

**Figure 5.**
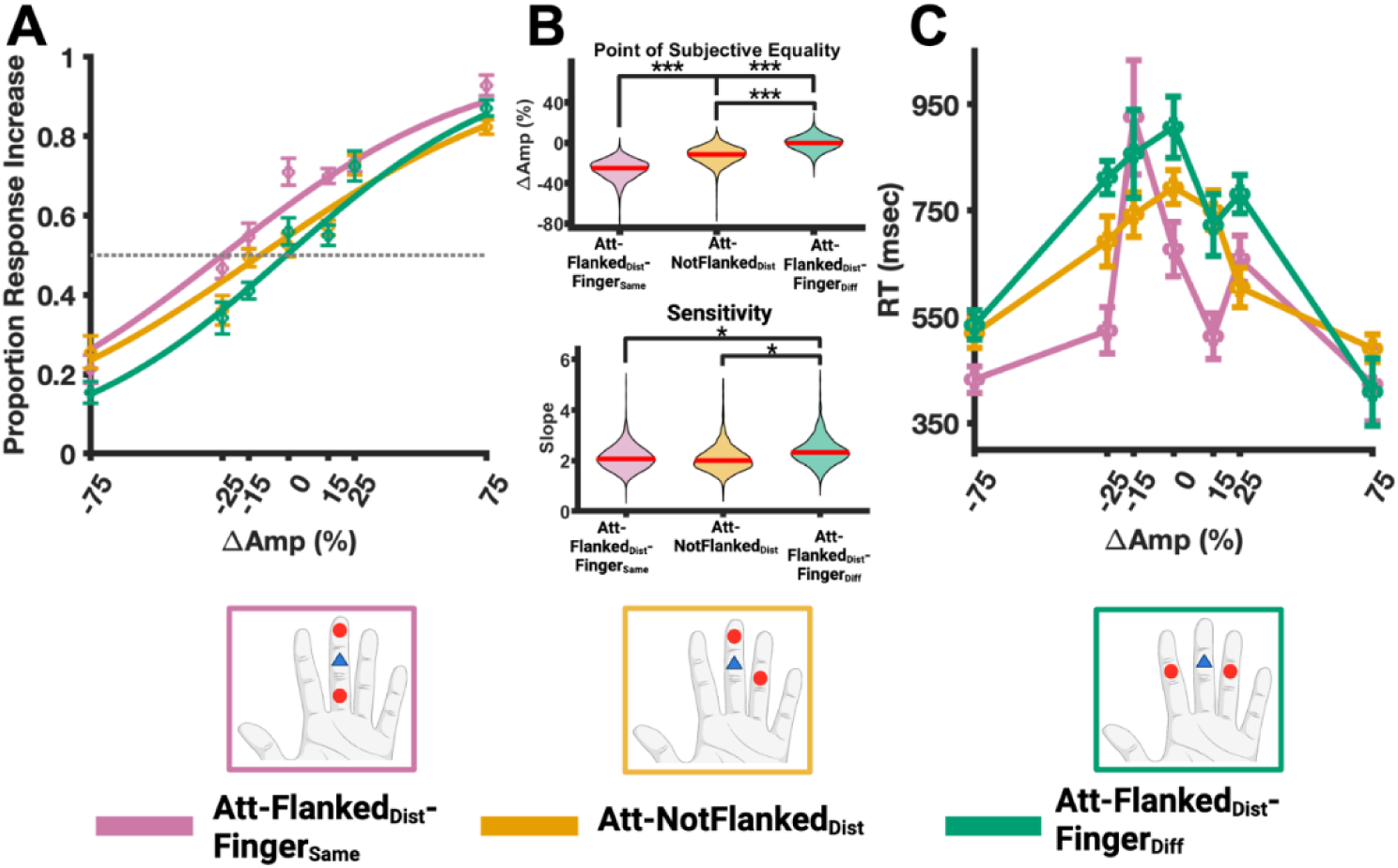
Two-Distractor spatial arrangement effects on perceptual sensitivity, response bias, and reaction time. (**A**) Psychometric fit (using logistic function) of the median responses across participants for a trial with 2 distractors on the same finger (magenta trace), different fingers and flanking the attended stimulus (orange trace), and different fingers but not flanking the attended stimulus (green trace). (**B**) The upper and lower graphs show violin plots of the PSE and perceptual sensitivity values of the psychometric function in (A), respectively. (**C**) Median RTs for each stimulus amplitude level. * = p < 0.05, ** = p < 0.01, *** = p < 0.001. N = 10. Error bars indicate ± within-subject SEM. Att-Flanked_Dist_-Finger_Same_ = Attended Stimulus Flanked by Distractors on Same Finger, Att-Flanked_Dist_-Finger_Diff_ = Attended Stimulus Flanked by Distractors on Different Fingers, and Att-NotFlanked_Dist_ = Attended Stimulus Not Flanked by Distractors.

The data also showed that RT is affected by the spatial arrangement of distractors (**Figure 5C**). A GLME model with factors of Stimulus Amplitude Level and Distractor Spatial Arrangement shows that the spatial arrangement of distractors has a significant effect on RT (F(2,176) = 3.1139, p = 0.04688), but Stimulus Amplitude Level does not have an effect on RT (F(1,176) = 1.852, p = 0.1753). The GLME model also did not show an interaction effect between Distractor Spatial Arrangement and Stimulus Amplitude Level (F(2,176) = 0.5501, p = 0.5779). The RT plot in **Figure 5C** indicates that there is a speed-accuracy tradeoff across the different Spatial Arrangement conditions as RT tends to be slower in the Att-Flanked_Dist_-Finger_Diff_ condition and faster in the Att-Flanked_Dist_-Finger_Same_ condition.

### Effects of cueing attended vs. distractor stimuli

**Figure 6A** show psychometric curves for valid (high probability of occurrence) and invalid (low probability of occurrence) conditions when attended stimuli (blue) or distractor stimuli (red) are cued. Non-parametric shuffling tests show no significant differences between validly- and invalidly-cued attended (p = 0.3270) nor distractor stimuli (p = 0.1212; *top panel* of **Figure 6B**). However, non-parametric shuffling tests demonstrate a significant difference in perceptual sensitivity for validly- vs. invalidly-cued attended stimuli (p = 0), but not for distractor stimuli (p = 0.1502; *bottom panel* of **Figure 6B**).

**Figure 6.**
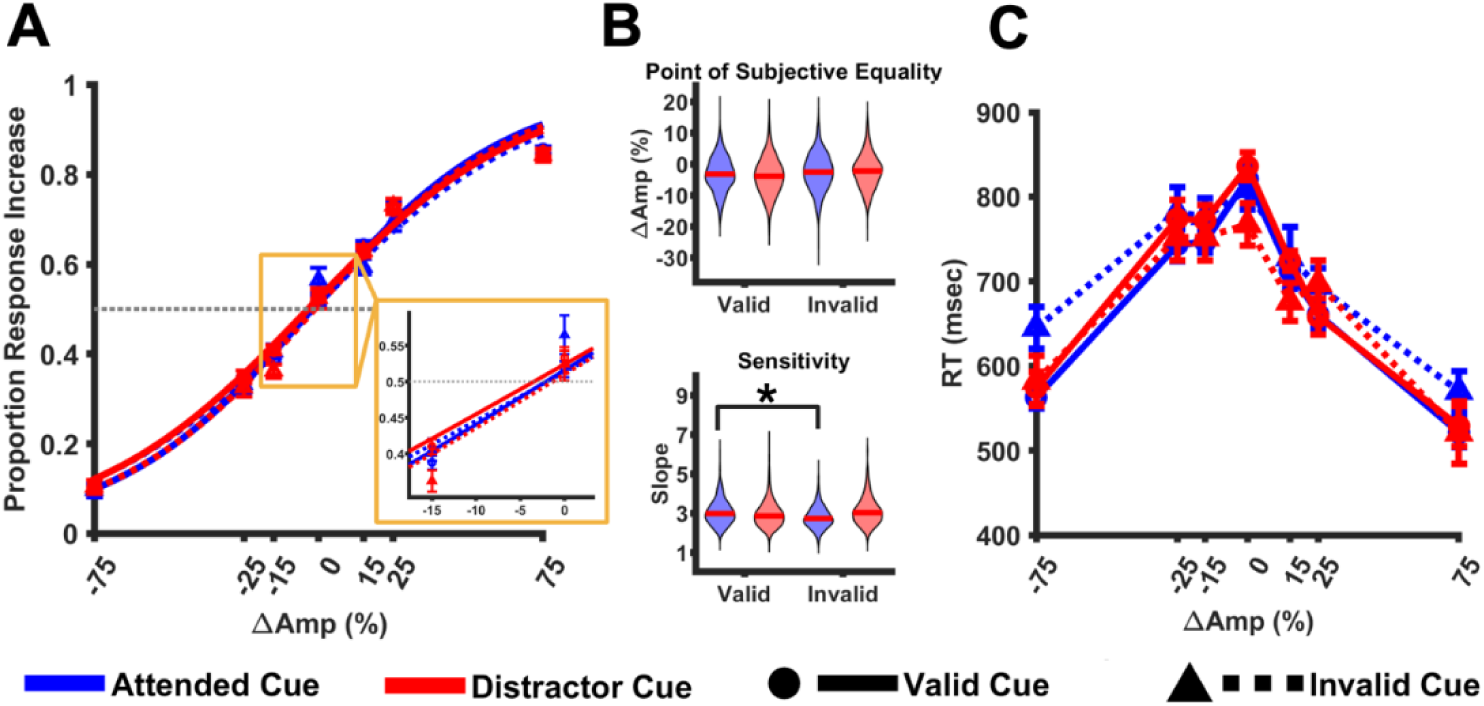
Cue type and validity effects on perceptual bias, sensitivity, and RT. (**A**) The left graph shows the psychometric functions for validly (solid traces) and invalidly (dotted traces) cueing the attended stimulus (blue traces) and distractor (red traces) locations. The solid lines are curve fits to the human data using a logistic function. The curve fits produced R^2^ values of 0.79, 0.72, 0.77, and 0.70 for the Valid-Attended, Invalid-Attended, Valid-Distractor, and Invalid-Distractor cue conditions, respectively. **(B)** Violin plots of the psychometric fit parameters, Point of subjective equality (PSE, middle-top) and Perceptual Sensitivity (middle-bottom). The solid red lines are the median values of each distribution, and the dotted black lines are the linear fits as a function of number of distractors. **(C)** The right graph shows the median RTs for each stimulus amplitude level. * = p < 0.05, N = 20. Error bars indicate ± within-subject SEM.

Cue Validity of attended and distractor stimuli also have small effect on RT (**Figure 6C**). A GLME model on RT with factors of Stimulus Amplitude Level, Cue Type, and Cue Validity show a trend towards significance effect of Stimulus Amplitude Level (F(1,552) = 3.1225, p = 0.0778). The GLME did not show a main effect of Cue Type (F(1,552) = 0.1706, p = 6798) or Cue Validity (F(1,552) = 0.0733, p = 0.7866). The GLME also did not show interaction effects between Stimulus Amplitude Level and Cue Type (F(1,552) = 0.0021, p = 0.9636), Stimulus Amplitude Level and Cue Validity (F(1,552) = 0.0230, p = 0.8796), Cue Type and Cue Validity (F(1,552) = 0.8825, p = 0.3479), nor a three-way interaction between Stimulus Amplitude Level, Cue Type, and Cue Validity (F(1,552) = 0.0052, p = 0.9427).

### Effects of attended vs. distractor cue spatial arrangement on performance

To address whether selective attention or distractor suppression utilizes a single vs. divided spotlight mechanism, we analyzed validity effects of attended and distractor cued conditions for stimuli presented in different spatial arrangements in Experiment 2. We first aimed to replicate the findings by (Eimer & Forster, 2003b) that showed that the attentional spotlight can be flexibly split across non-contiguous finger locations. Specifically, (Eimer & Forster, 2003b) found that attention effects on neural data were unaffected when a distractor location (e.g., digit 3) was between two possible attended locations on the hand (e.g., digits 2 and 5). In our experiment, we compared behavior between Distractor Flanked by Attended Stimuli on Different Fingers (i.e., distractors presented in the middle pad of digit 3 and attended stimuli presented in the middle pads of digits 2 and 4) vs. Distractor Not-Flanked by Attended Stimuli (e.g., distractors presented in the middle pad of digit 3 and attended stimuli presented in the distal pad of digit 3 and middle pad of digit 4) to determine whether the attentional spotlight can be split across non-contiguous finger locations. Consistent with previous observations on the attentional spotlight (Eimer & Forster, 2003b), a GLME model with factors of Spatial Arrangement (Distractor Flanked by Attended Stimuli on Different Fingers, Dist-Flanked-_Att_-Finger_Diff,_ and Distractor Not-Flanked by Attended Stimuli, Dist-NotFlanked_Att_) and Cue Validity (Valid vs. Invalid) did not reveal a main effect of Spatial Arrangement (F(1,61) = 0.07964, p = 0.7787), Cue Validity (F(1,61) = 0.4049, p = 0.5270), or an interaction effect (F(1,61) = 15306, p = 0.2208; *right panel* **Figure 7A**).

**Figure 7.**
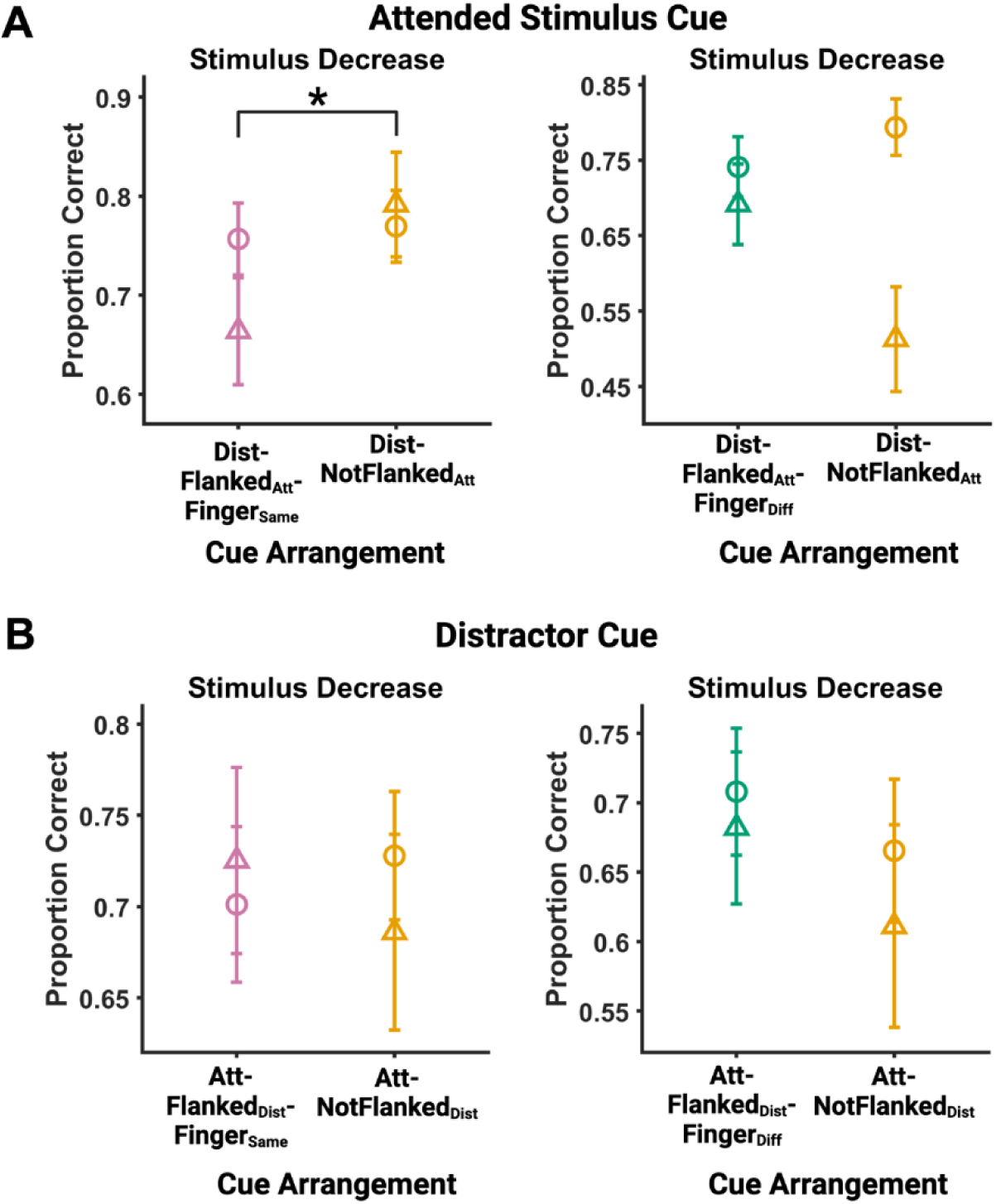
Spatial arrangement effects on cueing attended stimulus vs. distractor locations. (**A**) The graphs show the median accuracy across participants for stimuli with a decrease amplitude change for conditions where the attended stimulus locations are cued in a Same Finger Flanking vs. Mixed Finger Non-flanking arrangement (left) and Different Finger Flanking vs. Mixed Finger Non-flanking arrangement (right). Circles and triangles indicate when the attended stimulus occurs at the valid cue location and invalid cue location, respectively. (**B**) Same as (A) but for when the distractor locations are cued in a Same Finger Flanking vs. Mixed Finger Non-flanking arrangement (left) and Different Finger Flanking vs. Mixed Finger Non-flanking arrangement (right). * = p < 0.05, N = 20. Error bars indicate ± within-subject SEM. Att-Flanked_Dist_-Finger_Same_ = Attended Stimulus Flanked by Distractors on Same Finger, Att-Flanked_Dist_-Finger_Diff_ = Attended Stimulus Flanked by Distractors on Different Finger, and Att-NotFlanked_Dist_ = Attended Stimulus Not Flanked by Distractors, DistFlanked_Att_-Finger_Same_ = Distractor Flanked by Attended Stimuli on Same Finger, Dist-Flanked_Att_-Finger_Diff_ = Distractor Flanked by Attended Stimuli on Different Fingers, and Dist-NotFlanked_Att_ = Distractor Not Flanked by Attended Stimuli.

We next determined whether the spotlight of attention can be flexibly allocated to non-contiguous locations within the same finger. A GLME model with factors of Spatial Arrangement (Distractor Flanked by Attended Stimuli on Same Finger, Dist-Flanked_Att_-Finger_Same,_ vs. Distractor Not-Flanked by Attended Stimuli, Dist-NotFlanked_Att_) and Cue Validity (Valid vs. Invalid) showed that there is a main effect of Spatial Arrangement (F(1,64) = 5.4346, p = 0.0229), with performance impaired when attended stimuli are flanked by distractors within the same finger (*left panel* **Figure 7A**). The GLME did not reveal a main effect of Cue Validity (F(1,64) = 0.0002, p = 0.9890), but we did observe an interaction effect between Spatial Arrangement and Cue Validity (F(1,64) = 4.4039, p = 0.0398), with performance accuracy being lower when the attended stimulus occurred in the invalidly cued location in the same finger (magenta points, *left panel* **Figure 7A**). Finally, we also did not observe Spatial Arrangement and Validity effects on accuracy for amplitude increases or RT (see **Supplemental Figure 3**). Taken together, our results are highly consistent with those from (Eimer & Forster, 2003b) showing that the attentional spotlight can be allocated to non-contiguous areas across different fingers, but is more difficult to divide across non-contiguous phalanges within the same finger.

A major goal of this experiment is to determine whether the spotlight of distractor suppression can also be flexibly split across non-contiguous skin locations. To test this hypothesis, we compared behavior between Attended Stimuli Flanked by Distractors on Different Fingers (i.e., attended stimuli presented in the middle pad of digit 3 and distractors presented in the middle pads of digits 2 and 4) vs. Attended Stimuli Not-Flanked by Distractors (e.g., distractors presented in the middle pad of digit 3 and attended stimuli presented in the distal pad of digit 3 and middle pad of digit 4). Similar to our findings (*right panel* **Figure 7A**) and (Eimer & Forster, 2003b), a GLME test with factors of Spatial Arrangement (Attended Stimuli Flanked by Distractors on Different Fingers, Att-Flanked_Dist_-Finger_Diff_, vs. Attended Stimuli Not-Flanked by Distractors, Att-NotFlanked_Dist_) and Cue Validity (Valid vs. Invalid) did not show significant effects of Spatial Arrangement (F(1,64) = 0.5058, p = 0.4796), Cue Validity (F(1,64) = 1.2131, p = 0.2749), or an interaction effect (F(1,64) = 0.7045, p = 0.4044). Further, a GLME model with factors of Spatial Arrangement (Attended Stimuli Flanked by Distractors on the Same Finger, Att-Flanked_Dist_-Finger_Same_ vs. Attended Stimuli Not-Flanked by Distractors, Att-NotFlanked_Dist_) and Cue Validity (Valid vs. Invalid) did not show effects of Spatial Arrangement (F(1,65) = 0.06659, p = 0.7972), Cue Validity (1,65) = 1.1343, p = 0.2908), or an interaction effect (F(1,65) = 0.6202, p = 0.4338). These results suggest that distractor suppression mechanisms can be flexibly allocated to non-contiguous finger locations to facilitate vibration discrimination of attended stimuli.

## DISCUSSION

This study investigated the spatial flexibility properties of distractor suppression mechanisms in the tactile modality. Participants performed a series of vibration discrimination tasks on the hand with distractor stimuli presented in different spatial arrangements relative to the behaviorally-relevant stimulus. Our data show that cueing the location of distractors provides a perceptual benefit in discriminating behaviorally-relevant stimuli. Specifically, we observed that performance is enhanced for validly- vs. invalidly-cued stimuli (**Figures 3A and 3B**), and that invalidly cued distractors shift perceptual thresholds in favor of reporting stimulus amplitude increases (**Figure 2B**). The importance of cueing the distractor location is further evidenced by the substantial impairment in behavior when presenting a distractor when the cue indicated that no distractors would be presented (valid vs. invalid 0 distractor conditions; **Figures 3A and 3B** *gray bars*). Taken together, these data show that proactive deployment of distractor suppression enhances perception of attended stimuli, an effect that is also observed in the visual system (Redding & Fiebelkorn, 2024a).

Distractor interference effects vary depending on the spatial arrangement of the distractors, with performance decreasing when distractors are presented within the same finger vs. different fingers relative to attended stimuli (**Figures 5 and 7A**). These data suggest that the spatial resolution of tactile selective attention might be limited to a finger, a finding that aligns with previous observations in evoked related potentials (ERPs) in humans (Eimer & Forster, 2003b). The study by Eimer and Forster (2003b) showed that neural responses to the cue and attended stimuli presented on the same finger location were enhanced as compared to when the cue and attended stimuli were presented on different phalanxes of the same finger (e.g., cue and stimulus on the middle finger pad of the index finger vs. cue and stimulus on the middle and proximal finger pad of the index finger, respectively). However, neural activity to the latter condition was greater than activity in response to when the cue and stimuli were presented on different fingers (e.g., cue on the middle finger pad of the index finger and the stimulus presented on the proximal finger pad of the middle finger). Of note, (Eimer & Forster, 2003b) did not present distractor stimuli (i.e., stimuli that were instructed to be ignored), but rather used cueing probability to determine the expected location of the attended stimuli. Our experiment, however, used distractor stimuli and cued the location of the distractors on a trial-by-trial basis. Thus, it seems that both mechanisms of attentional enhancement and distracter suppression are spatially restricted to a single finger.

Our data provide evidence that the spotlight of suppression can be split across non-contiguous skin locations. When cueing distractors, we observed that behavior is unaltered when distractors are presented across non-neighboring areas (**Figure 7B** *lower graphs*). Similarly, when cueing attended stimuli, our data show that behavior is unaffected when attended stimuli are presented across non-neighboring fingers, a finding that is consistent with previous studies of attention in touch (Eimer & Forster, 2003b). However, our study showed that behavior is impaired when attended stimuli are presented across non-neighboring phalanges within a finger (**Figure 7B** *upper panels*). These data suggest that, while the spotlight of attention can be flexibly split across fingers but limited within a finger, mechanisms of suppression might be more flexible such that the spotlight of suppression can be split within and across fingers. Neural recordings, especially at the single cell level, should be conducted to further validate this hypothesis. However, the different pattern of effects between cueing attended vs. distractor stimuli strongly support the general hypothesis that mechanisms of suppression are separable than those used for enhancing behaviorally-relevant inputs, a finding that is supported by visual studies (Chelazzi et al., 2019; Redding & Fiebelkorn, 2024b)

Our findings show that tactile distractors cannot be fully ignored, with distractor effects occurring for stimuli with reductions in amplitude vibration only. In particular, we found that perceptual threshold and perceptual sensitivity are systematically impaired with the number of distractors (**Figure 2A**). These results align with previous literature showing that as more distractors are presented, the harder it is to perceive a target tactile stimulus (Gherri, Fiorino, et al., 2023; Halfen et al., 2020). These previous studies demonstrate distractor effects in the context of tactile search, where the location of the distractors is not specified, and thus not proactively suppressed. In our study, we show that even if proactive suppression can be deployed, tactile distractors cannot be fully ignored and still affect the ability to discriminate attended tactile stimuli. We also observe that distractor effects are dependent on participants’ perceptual biases, with accuracy and RTs scaling as a function of PSE_PreTask_ (**Figure 4**). Specifically, the data showed that participants with lower PSEs (e.g., negative) have lower accuracy and higher RTs as compared to participants with higher PSEs. Negative and positive PSE_PreTask_ values indicate that participants have a bias in reporting that stimuli increase and decrease in amplitude, respectively. Thus, it seems that participants with more negative biases are more susceptible to distractor interference effects, such that stimuli that decrease in amplitude are judged as increasing in amplitude. Notably, the PSE_PreTask_ and behavior relationship is not due to participants’ perceptual abilities because the relationship does not occur when distractors are not presented (i.e., in 0 Distractor conditions) and the relationship scales as a function of distractor number (see **Figure 4**).

In conclusion, our results show that proactive deployment of distractor suppression aids perception, and that mechanisms of distractor suppression are separable from mechanisms used for enhancing attended stimuli. The results also show that distractor interference effects depend on the spatial arrangement of the distractors relative to attended stimuli, with stronger interference for distractors placed in the same finger relative to the attended stimuli. Further, our data provide evidence that the spotlight of distractor suppression can be split across non-contiguous areas within and across fingers. We stress that these conclusions are based on behavioral data, and need to be validated with neurophysiological evidence, particularly at the single cell level.

---

**Supplemental Figure 1:**
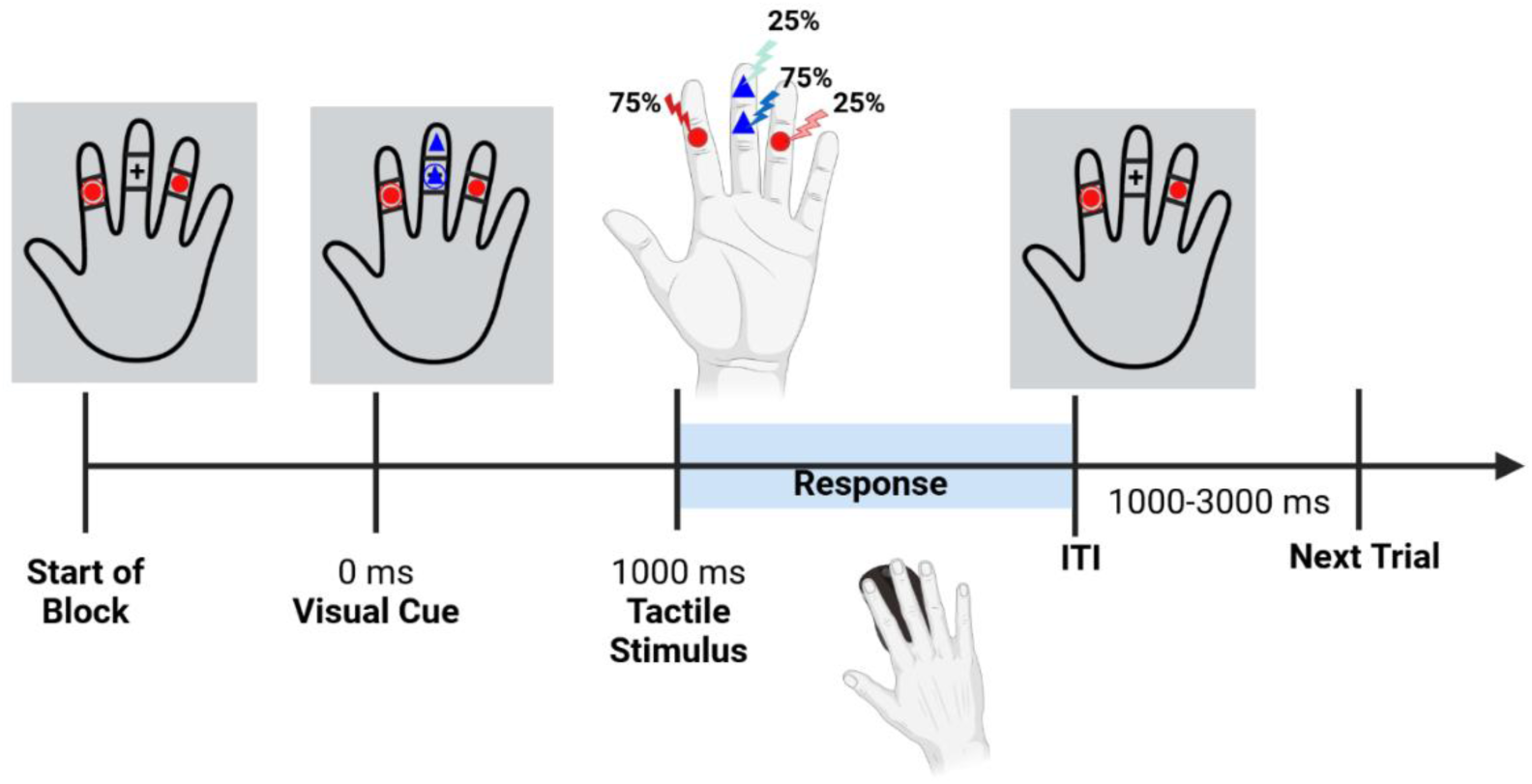
Sequence of events in a trial of the amplitude discrimination task in Experiment 2. In each block of trials, either the two attended or distractor locations were fixed, while the locations of the other cue type were flexibly cued on a trial-by-trial basis. Red circles indicated the distractor locations, while blue triangles indicated the attended locations. The cues of the fixed locations remained on the VR screen throughout the entirety of the block. Each trial began with the visual cue showing the locations of the flexible cue type. After 1000 ms, the target and distractor vibrations were provided simultaneously. For each cue type, the stimulus occurred at the cued location with the ring around it for 75% of trials (i.e., high-probability or valid location), and occurred at the other cued location for the other 25% of trials (i.e., low-probability or invalid location). The target vibration started at a baseline amplitude (subject’s PSE), then increased or decreased in amplitude after 500 ms for another 500 ms. The distractor vibration remained at a constant amplitude, which was set at half of the baseline amplitude of the target vibration. Participants made their responses with mouse button clicks.

**Supplemental Figure 2:**
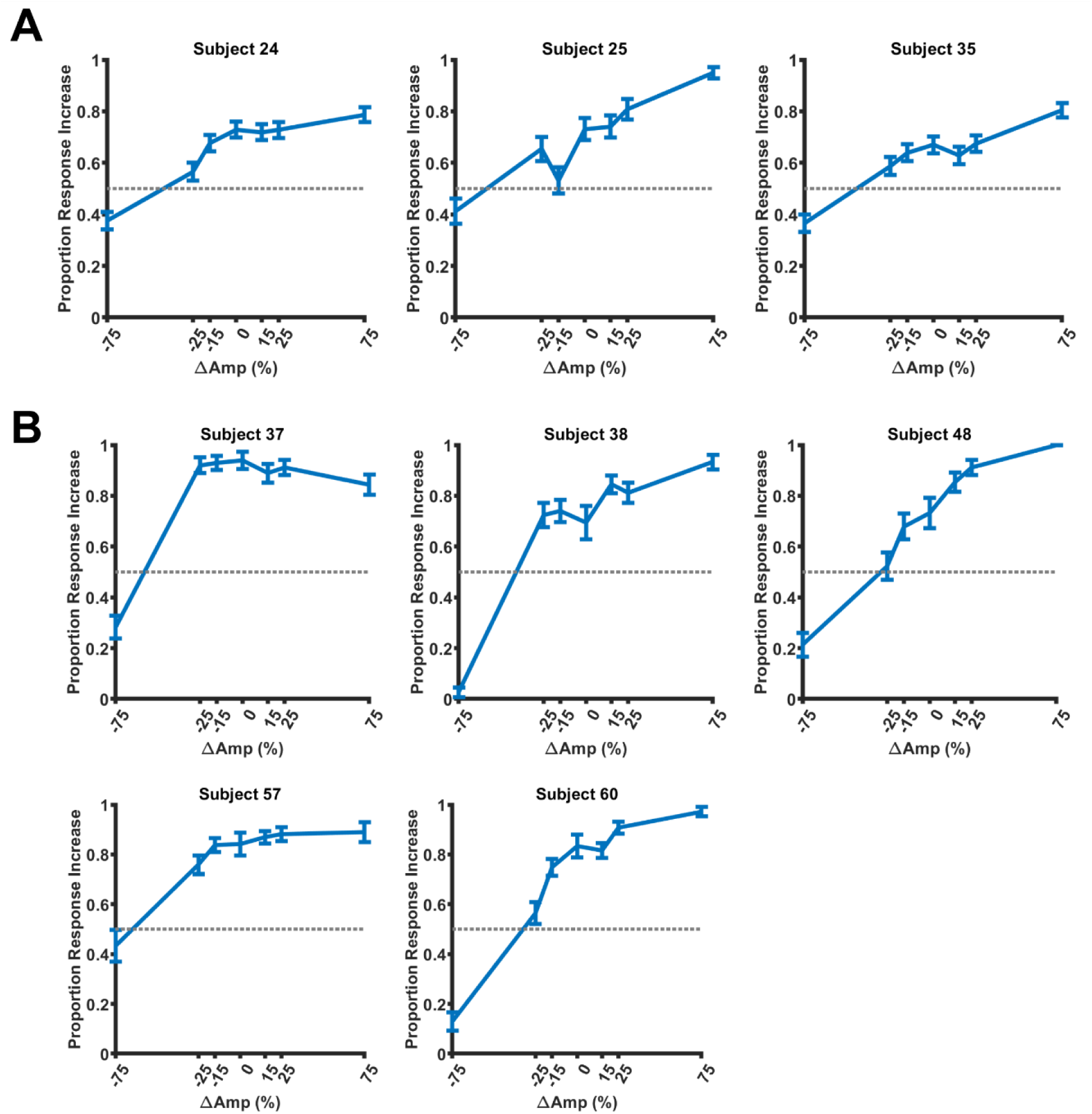
Psychometric functions of participants that were not included in the analyses for (**A**) Experiment 1 and (**B**) Experiment 2. These psychometric curves illustrate participants with a strong bias towards indicating an amplitude increase in the stimulus. These participants were removed because five or more points in the stimulus amplitude change condition elicited significant responses above 0.5 proportion rate.

**Supplemental Figure 3.**
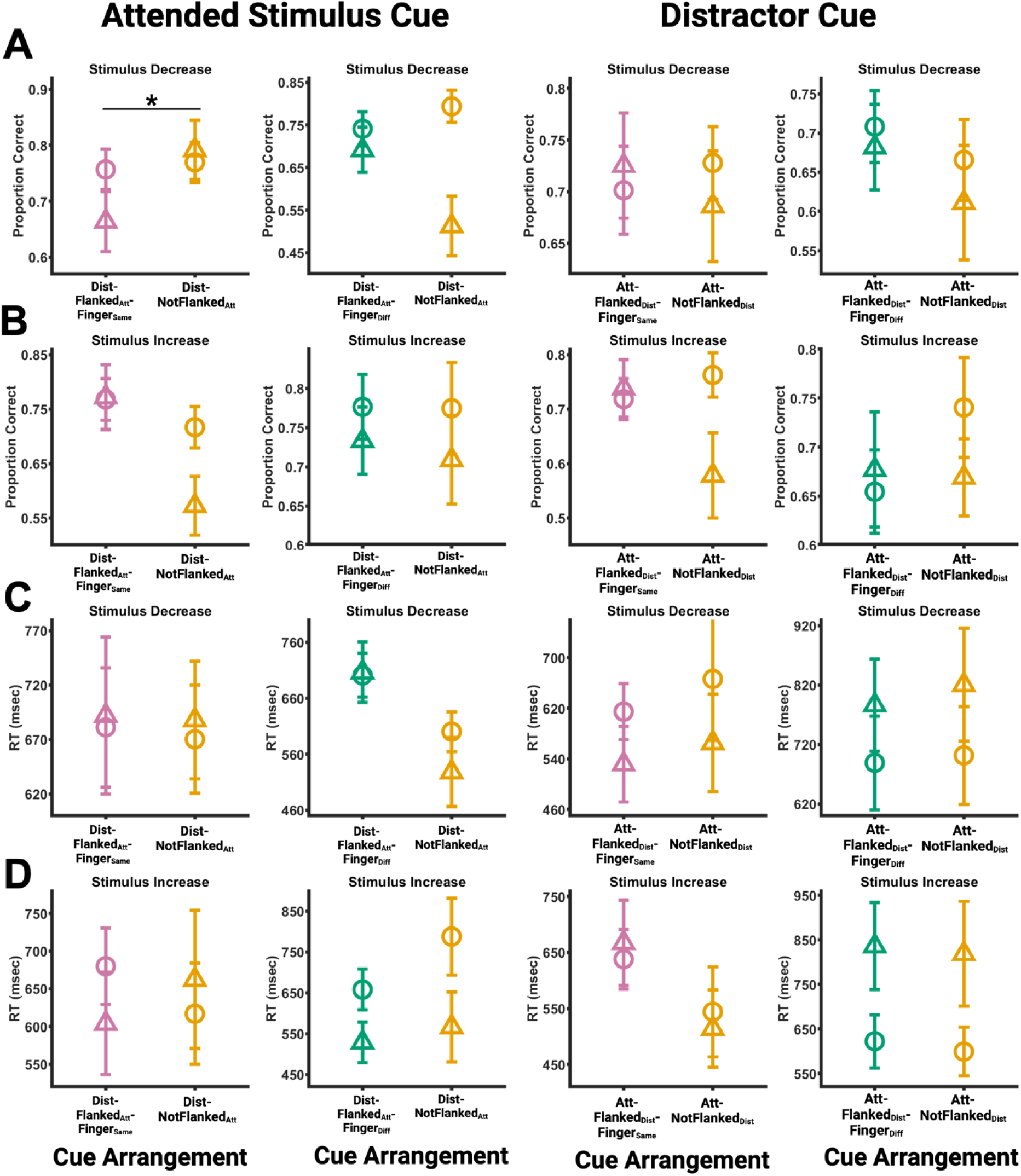
Spatial arrangement effects on cueing attended stimulus vs. distractor locations. (**A**) The graphs show the median accuracy across participants for stimuli with a decrease amplitude change when attended stimuli are cued in the Same Finger Flanking vs. Mixed Finger Non-flanking (far left) and Different Finger Flanking vs. Mixed Finger Non-flanking (middle right) arrangements and when distractors are cued in the Same Finger Flanking vs. Mixed Finger Non-flanking (middle right) and Different Finger Flanking vs. Mixed Finger Non-flanking (far right) arrangements. Circles and triangles indicate when the attended stimulus occurs at the valid cue location and invalid cue location, respectively. (**B**) Same as (A) but for median accuracy across participants for stimuli with increase amplitude changes. (**C**) Same as (A) but for median RT across participants for stimuli with decrease amplitude changes. (**D**) Same as (A) but for median RT across participants for stimuli with increase amplitude changes. * = p < 0.05, N = 20. Error bars indicate ± within-subject SEM. Att-Flanked_Dist_-Finger_Same_ = Attended Stimulus Flanked by Distractors on Same Finger, Att-Flanked_Dist_-Finger_Diff_ = Attended Stimulus Flanked by Distractors on Different Finger, and Att-NotFlanked_Dist_ = Attended Stimulus Not Flanked by Distractors, DistFlanked_Att_-Finger_Same_ = Distractor Flanked by Attended Stimuli on Same Finger, Dist-Flanked_Att_-Finger_Diff_ = Distractor Flanked by Attended Stimuli on Different Fingers, and Dist-NotFlanked_Att_ = Distractor Not Flanked by Attended Stimuli.

## Notes

### Competing Interest Statement

The authors have declared no competing interest.

## REFERENCES

Adler, J., Giabbiconi, C.-M., & Müller, M. M. (2009). Shift of attention to the body location of distracters is mediated by perceptual load in sustained somatosensory attention. Biological Psychology, 81(2), 77–85. 10.1016/j.biopsycho.2009.02.001

Braithwaite, J. J., Hulleman, J., Andrews, L., & Humphreys, G. W. (2010). Measuring the spread of spreading suppression: A time-course analysis of spreading suppression and its impact on attentional selection. Vision Research, 50(3), 346–356. 10.1016/j.visres.2009.11.019

Chang, S., & Egeth, H. E. (2019). Enhancement and Suppression Flexibly Guide Attention. Psychological Science, 30(12), 1724–1732. 10.1177/0956797619878813

Chelazzi, L., Marini, F., Pascucci, D., & Turatto, M. (2019). Getting rid of visual distractors: The why, when, how, and where. Current Opinion in Psychology, 29, 135–147. 10.1016/j.copsyc.2019.02.004

Clepper, G. M., Martinez, J. S., & Tan, H. Z. (2021). Selective and Divided Attention for Vibrotactile Stimuli on Both Arms. 2021 IEEE World Haptics Conference (WHC), 650–655. 10.1109/WHC49131.2021.9517140

Craig, J. C. (1985). Attending to two fingers: Two hands are better than one. Perception & Psychophysics, 38(6), 496–511. 10.3758/BF03207059

Daniel, S., Andrillon, T., Tsuchiya, N., & van Boxtel, J. J. A. (2022). Divided attention in the tactile modality. *Attention, Perception*, & Psychophysics, 84(1), 47–63. 10.3758/s13414-021-02352-8

Drisdelle, B. L., & Eimer, M. (2023). Proactive suppression can be applied to multiple salient distractors in visual search. Journal of Experimental Psychology: General, 152(9), 2504–2519. 10.1037/xge0001398

Ede, F. van, Lange, F. de, Jensen, O., & Maris, E. (2011). Orienting Attention to an Upcoming Tactile Event Involves a Spatially and Temporally Specific Modulation of Sensorimotor Alpha- and Beta-Band Oscillations. Journal of Neuroscience, 31(6), 2016–2024. 10.1523/JNEUROSCI.5630-10.2011

Eimer, M., & Forster, B. (2003a). Modulations of early somatosensory ERP components by transient and sustained spatial attention. Experimental Brain Research, 151(1), 24–31. 10.1007/s00221-003-1437-1

Eimer, M., & Forster, B. (2003b). The spatial distribution of attentional selectivity in touch: Evidence from somatosensory ERP components. Clinical Neurophysiology, 114(7), 1298–1306. 10.1016/S1388-2457(03)00107-X

Evans, P. M., & Craig, J. C. (1991). Tactile attention and the perception of moving tactile stimuli. Perception & Psychophysics, 49(4), 355–364. 10.3758/bf03205993

Forster, B., & Eimer, M. (2005). Covert attention in touch: Behavioral and ERP evidence for costs and benefits. Psychophysiology, 42(2), 171–179. 10.1111/j.1469-8986.2005.00268.x

Forster, B., Tziraki, M., & Jones, A. (2016). The attentive homunculus: ERP evidence for somatotopic allocation of attention in tactile search. Neuropsychologia, 84, 158–166. 10.1016/j.neuropsychologia.2016.02.009

Foxe, J. J., Simpson, G. V., & Ahlfors, S. P. (1998). Parieto-occipital ∼1 0Hz activity reflects anticipatory state of visual attention mechanisms. NeuroReport, 9(17), 3929.

Foxe, J. J., & Snyder, A. C. (2011). The Role of Alpha-Band Brain Oscillations as a Sensory Suppression Mechanism during Selective Attention. Frontiers in Psychology, 2. 10.3389/fpsyg.2011.00154

Foxe, J., & Snyder, A. (2011). The Role of Alpha-Band Brain Oscillations as a Sensory Suppression Mechanism during Selective Attention. Frontiers in Psychology, 2. https://www.frontiersin.org/articles/10.3389/fpsyg.2011.00154

Frey, H.-P., Schmid, A. M., Murphy, J. W., Molholm, S., Lalor, E. C., & Foxe, J. J. (2014). Modulation of early cortical processing during divided attention to non-contiguous locations. The European Journal of Neuroscience, 39(9), 1499–1507. 10.1111/ejn.12523

Frings, C., Bader, R., & Spence, C. (2008). Selection in touch: Negative priming with tactile stimuli. Perception & Psychophysics, 70(3), 516–523. 10.3758/pp.70.3.516

Gaspelin, N., Leonard, C. J., & Luck, S. J. (2015). Direct Evidence for Active Suppression of Salient-but-Irrelevant Sensory Inputs. Psychological Science, 26(11), 1740–1750. 10.1177/0956797615597913

Gaspelin, N., & Luck, S. J. (2018). The Role of Inhibition in Avoiding Distraction by Salient Stimuli. Trends in Cognitive Sciences, 22(1), 79–92. 10.1016/j.tics.2017.11.001

Geng, J. J. (2014). Attentional Mechanisms of Distractor Suppression. Current Directions in Psychological Science, 23(2), 147–153. 10.1177/0963721414525780

Gherri, E., White, F., & Venables, E. (2023). On the spread of spatial attention in touch: Evidence from event-related brain potentials. Biological Psychology, 178, 108544. 10.1016/j.biopsycho.2023.108544

Gomez-Ramirez, M., Hysaj, K., & Niebur, E. (2016a). Neural mechanisms of selective attention in the somatosensory system. Journal of Neurophysiology, 116(3), 1218–1231. 10.1152/jn.00637.2015

Gomez-Ramirez, M., Hysaj, K., & Niebur, E. (2016b). Neural mechanisms of selective attention in the somatosensory system. Journal of Neurophysiology, 116(3), 1218–1231. 10.1152/jn.00637.2015

Haegens, S., Luther, L., & Jensen, O. (2012a). Somatosensory Anticipatory Alpha Activity Increases to Suppress Distracting Input. Journal of Cognitive Neuroscience, 24(3), 677–685. 10.1162/jocn_a_00164

Haegens, S., Luther, L., & Jensen, O. (2012b). Somatosensory Anticipatory Alpha Activity Increases to Suppress Distracting Input. Journal of Cognitive Neuroscience, 24(3), 677–685. 10.1162/jocn_a_00164

Hsiao, S. S., & Gomez-Ramirez, M. (2012). Neural Mechanisms of Tactile Perception. In Handbook of Psychology, *Second Edition*. John Wiley & Sons, Ltd. 10.1002/9781118133880.hop203008

Hughes, R. W. (2014). Auditory distraction: A duplex-mechanism account. PsyCh Journal, 3(1), 30–41. 10.1002/pchj.44

Jans, B., Peters, J. C., & De Weerd, P. (2010). Visual spatial attention to multiple locations at once: The jury is still out. Psychological Review, 117(2), 637–684. 10.1037/a0019082

Jones, S. R., Kerr, C. E., Wan, Q., Pritchett, D. L., Hämäläinen, M., & Moore, C. I. (2010). Cued Spatial Attention Drives Functionally Relevant Modulation of the Mu Rhythm in Primary Somatosensory Cortex. Journal of Neuroscience, 30(41), 13760–13765. 10.1523/JNEUROSCI.2969-10.2010

Kelly, S. P., Foxe, J. J., Newman, G., & Edelman, J. A. (2010). Prepare for conflict: EEG correlates of the anticipation of target competition during overt and covert shifts of visual attention. European Journal of Neuroscience, 31(9), 1690–1700. 10.1111/j.1460-9568.2010.07219.x

Kelly, S. P., Gomez-Ramirez, M., & Foxe, J. J. (2008). Spatial Attention Modulates Initial Afferent Activity in Human Primary Visual Cortex. Cerebral Cortex, 18(11), 2629– 2636. 10.1093/cercor/bhn022

McMains, S. A., & Somers, D. C. (2004). Multiple Spotlights of Attentional Selection in Human Visual Cortex. Neuron, 42(4), 677–686. 10.1016/S0896-6273(04)00263-6

Mohr, K. S., Carr, N., Georgel, R., & Kelly, S. P. (2020). Modulation of the Earliest Component of the Human VEP by Spatial Attention: An Investigation of Task Demands. Cerebral Cortex Communications, 1(1), tgaa045. 10.1093/texcom/tgaa045

Moore, T., & Zirnsak, M. (2017). Neural Mechanisms of Selective Visual Attention. Annual Review of Psychology, 68(Volume 68, 2017), 47–72. 10.1146/annurev-psych-122414-033400

Noonan, M. P., Adamian, N., Pike, A., Printzlau, F., Crittenden, B. M., & Stokes, M. G. (2016). Distinct Mechanisms for Distractor Suppression and Target Facilitation. Journal of Neuroscience, 36(6), 1797–1807. 10.1523/JNEUROSCI.2133-15.2016

Noonan, M. P., Crittenden, B. M., Jensen, O., & Stokes, M. G. (2018). Selective inhibition of distracting input. Behavioural Brain Research, 355, 36–47. 10.1016/j.bbr.2017.10.010

Redding, Z. V., & Fiebelkorn, I. C. (2024a). Separate Cue- and Alpha-Related Mechanisms for Distractor Suppression. Journal of Neuroscience, 44(25). 10.1523/JNEUROSCI.1444-23.2024

Redding, Z. V., & Fiebelkorn, I. C. (2024b). Separate Cue- and Alpha-Related Mechanisms for Distractor Suppression. The Journal of Neuroscience, 44(25), e1444232024. 10.1523/JNEUROSCI.1444-23.2024

Sacchet, M. D., LaPlante, R. A., Wan, Q., Pritchett, D. L., Lee, A. K. C., Hämäläinen, M., Moore, C. I., Kerr, C. E., & Jones, S. R. (2015). Attention Drives Synchronization of Alpha and Beta Rhythms between Right Inferior Frontal and Primary Sensory Neocortex. Journal of Neuroscience, 35(5), 2074–2082. 10.1523/JNEUROSCI.1292-14.2015

Schneider, D., Herbst, S. K., Klatt, L.-I., & Wöstmann, M. (2022). Target enhancement or distractor suppression? Functionally distinct alpha oscillations form the basis of attention. European Journal of Neuroscience, 55(11–12), 3256–3265. 10.1111/ejn.15309

Schwartz, Z. P., & David, S. V. (2018). Focal Suppression of Distractor Sounds by Selective Attention in Auditory Cortex. Cerebral Cortex, 28(1), 323–339. 10.1093/cercor/bhx288

Snyder, A. C., & Foxe, J. J. (2010). Anticipatory attentional suppression of visual features indexed by oscillatory alpha-band power increases: A high-density electrical mapping study. The Journal of Neuroscience: The Official Journal of the Society for Neuroscience, 30(11), 4024–4032. 10.1523/JNEUROSCI.5684-09.2010

van Ede, F., Köster, M., & Maris, E. (2012). Beyond establishing involvement: Quantifying the contribution of anticipatory α- and β-band suppression to perceptual improvement with attention. Journal of Neurophysiology, 108(9), 2352–2362. 10.1152/jn.00347.2012

van Ede, F., Szebényi, S., & Maris, E. (2014a). Attentional modulations of somatosensory alpha, beta and gamma oscillations dissociate between anticipation and stimulus processing. NeuroImage, 97, 134–141. 10.1016/j.neuroimage.2014.04.047

van Ede, F., Szebényi, S., & Maris, E. (2014b). Attentional modulations of somatosensory alpha, beta and gamma oscillations dissociate between anticipation and stimulus processing. NeuroImage, 97, 134–141. 10.1016/j.neuroimage.2014.04.047

van Moorselaar, D., & Slagter, H. A. (2020). Inhibition in selective attention. Annals of the New York Academy of Sciences, 1464(1), 204–221. 10.1111/nyas.14304

Vanrullen, R., & Dubois, J. (2011). The Psychophysics of Brain Rhythms. Frontiers in Psychology, 2. 10.3389/fpsyg.2011.00203

Worden, M. S., Foxe, J. J., Wang, N., & Simpson, G. V. (2000). Anticipatory Biasing of Visuospatial Attention Indexed by Retinotopically Specific α-Bank Electroencephalography Increases over Occipital Cortex. Journal of Neuroscience, 20(6), RC63–RC63. 10.1523/JNEUROSCI.20-06-j0002.2000

